# Bioinformatic analysis for the identification of potential gene interactions and therapeutic targets in atrial fibrillation

**DOI:** 10.1101/2020.05.18.101972

**Authors:** Shandong Yu, Jinyu Yu, Yong Guo, Yanpeng Chu, Heping Zhang

## Abstract

**Background:** Atrial fibrillation (AF) is the most prevalent tachycardia. The major injuries caused by AF are systemic embolism and heart failure. Although AF therapies have evolved substantially in recent years, the success rate of sinus rhythm maintenance is relatively low. The reason is the incomplete understanding of the AF mechanisms.

**Material and method:** In this study, profiles were downloaded from the GEO database (GSE79762). Bioinformatic analysis was used to identify differentially expressed genes (DEGs). GO analysis and KEGG analysis were performed to identify the most enriched terms and pathways. A protein-protein interaction network was constructed to determine regulatory genes. Key modules and hub genes were identified by MOCDE and cytoHubba. Transcription factors (TFs) were predicted by PASTAA.

**Results:** Seventy-seven up-regulated DEGs and 236 downregulated DEGs were identified. In the GO biological process, cellular components, and molecular function analysis, positive regulation of cell migration, anchoring junction and cell adhesion molecule binding were the most significant enrichment terms. The Hippo signaling pathway was the most significantly enriched pathway. In the PPI network analysis, we found that Class A/1 (rhodopsin-like receptors) may be the critical module in AF. Ten hub genes were extracted, including 4 upregulated genes and 6 downregulated genes. CXCR2, TLR4 and CXCR4 may play critical roles in AF. In TF prediction, we found that Irf-1 may be implicated in AF.

**Conclusion:** Our study found that the CXCR4, TLR4, CXCR2; Hippo signaling pathway; and class A/1 (rhodopsin-like receptors) modules may play critical roles in AF occurrence and maintenance. This may provide novel targets for AF treatment.

## 1. Introduction

Atrial fibrillation (AF) is the most prevalent cardiac arrhythmia. The incidence of AF is expected to double by 2030 because of the aging of the population. It is classified as paroxysmal AF, persistent AF and permanent AF[1]. The prevalence rate is more than 10% among those older than 80 years[2]. The major harms of AF are embolism (such as stroke) and heart failure. Effective treatment of AF will improve the clinical outcome of AF, such as reducing the disability rate caused by stroke. Although treatment strategies have evolved substantially in recent years, the efficacy is imperfect. Radiofrequency ablation of AF has evolved considerably to become safer and more effective over the past decade, while the recurrence rate is relatively high, especially for persistent AF[3]. Pharmacological treatment of AF may induce substantial adverse side effects, such as drug-induced proarrhythmia and cardiac and noncardiac toxicity[4–6]. The reason that current treatment strategies have limited efficacy is the incomplete understanding of the mechanisms of occurrence and progression of AF. Understanding the mechanisms may help us to find novel strategies for AF treatment. Successful cure of hepatitis C is due to a complete understanding of the mechanism of hepatitis C virus infection.

Omics has become increasingly important in investigating the mechanisms of diseases by comparing the differential expression of genes, RNAs or proteins between patients and controls. Since each study ultimately results in a list containing dozens or hundreds of gene candidates, it is essential to leverage existing biological knowledge through understanding the representation of known pathways or complexes within these datasets. Providing this molecular context can facilitate the interpretation of systems-level data and enable new discoveries[7]. Bioinformatic analysis is a powerful tool for omics dataset analysis. In this study, we used sample GSE79768 to analyze differentially expressed genes (DEGs) in patients with AF compared with patients with sinus rhythm. Functional enrichment analysis, such as gene ontology (GO) and Kyoto Encyclopedia of Genes and Genomes (KEGG) pathway analysis, and protein-protein interaction (PPI) analysis were performed to identify potential hub genes or key modules involved in AF.

## 2. Material and methods

### 2.1 Data source

The GSE79768 dataset (expression profiling generated by using GPL570 ([HG-U133_Plus_2] Affymetrix Human Genome U133 Plus 2.0 Array)[8] was downloaded from the GEO database (http://www.ncbi.nlm.nih.gov/geo/). In this dataset, paired left atrial and right atrial specimens (26 specimens) were obtained from 13 patients (7 persistent AF, 6 sinus rhythm). The specimens of atrial appendages were obtained from patients receiving surgery for mitral valve or coronary artery disease. Patients with AF presented persistent AF known more than 6 months, and patients with SR had no evidence of AF clinically without use of any anti-arrhythmic drugs.[2, 8].

### 2.2 Data processing

The robust multiarray analysis (RMA) method was used to process raw data of the dataset *(http://www.bioconductor.org)*. The probe ID for each gene was then converted to a gene symbol using hgu133a.db,org.Hs.eg.db and the annotate package in Bioconductor *(http://www.bioconductor.org)*[9].

### 2.3 Identification of DEGs

Differentially expressed genes (DEGs) in the left samples of patients with AF compared with those of patients with SR were identified by the R package LIMMA[10]. We used the false discovery rate (FDR) for p value correction by the Benjamin and Hochberg method[11] and for fold change (FC) calculation. Log2 FC < −1 / Log2 > 1 and adjusted p <0.05 were set as thresholds for DEGs.

### 2.4 Functional enrichment analysis

Gene Ontology (GO) and Kyoto Encyclopedia of Genes and Genomes pathway enrichment analyses were performed by using the database for annotation, visualization and integrated discovery bioinformatics resources (DAVID Gene Functional Classification Tool, http://david.abcc.ncifcrf.gov/)[12] and Metascape. The cutoff of the p value was 0.01. To further capture the relationships between the terms, a subset of enriched terms have been selected and rendered as a network plot, where terms with a similarity > 0.3 are connected by edges. We select the terms with the best p-values from each of the 20 clusters, with the constraint that there are no more than 15 terms per cluster and no more than 250 terms in total. The network is visualized using Cytoscape, where each node represents an enriched term and is colored first by its cluster ID and then by its p-value[7].

### 2.5 Construction of the differential coexpression gene network

To investigate interactions between proteins, a protein-protein interaction (PPI) network of differentially expressed genes was constructed by using the search tool for the retrieval of interacting genes (STRING database, V11.0; http://string-db.org/). The minimum required interaction score was 0.4[13].

### 2.6 Analysis of hub genes

The Cytohubba plugin of Cytoscape was used to determine key genes, also called hub genes, in the network. To obtain the key genes, we extracted the hub genes using cytoHubba. The MCC method was used to identify the top 10 hub genes[14].

### 2.7 Module

The Molecular Complex Detection (MCODE) algorithmhas been applied to identify densely connected network components. Pathway and process enrichment analysis was applied to each MCODE component independently, and the three best-scoring terms by p-value were retained[14].

### 2.8 Analysis of TFs

Transcription factors (TFs) were predicted by using PASTAA[15]. After DEGs were established, the calculated p value and association score were used to evaluate the correlation between the disease and TFs through hypergeometric distribution[9].

## 3. Results

### 3.1 The DEGs and Functional enrichment of DEGs

There were DEGs identified in left atrial specimens of patients with AF compared with patients with SR, including 77 upregulated DEGs and 236 downregulated DEGs. GO and KEGG pathway analyses were performed using the DAVID database and Metascape. The results are shown in Figure 2. The GO enrichment analysis and KEGG pathway analysis revealed that the identified DEGs were significant. The top 10 most significant GO terms of the enriched biological processes (BPs) (Figure 1A), cell components (CCs) (Figure 1B), and molecular function (MFs) (Figure 1C) are shown according to gene ratio. These include positive regulation of cell migration, anchoring junction and cell adhesion molecule binding. Moreover, the associated KEGG pathways most significantly enriched by DEGs are displayed in Figure 1D. The Hippo signaling pathway is the most significantly enriched pathway. The network of enriched terms colored by cluster ID is shown in Figure 2A, where nodes that share the same cluster ID are typically close to each other. Network colored by p-value is shown in Figure 2B, where terms containing more genes tend to have a more significant p-value.

**Figure 1.**
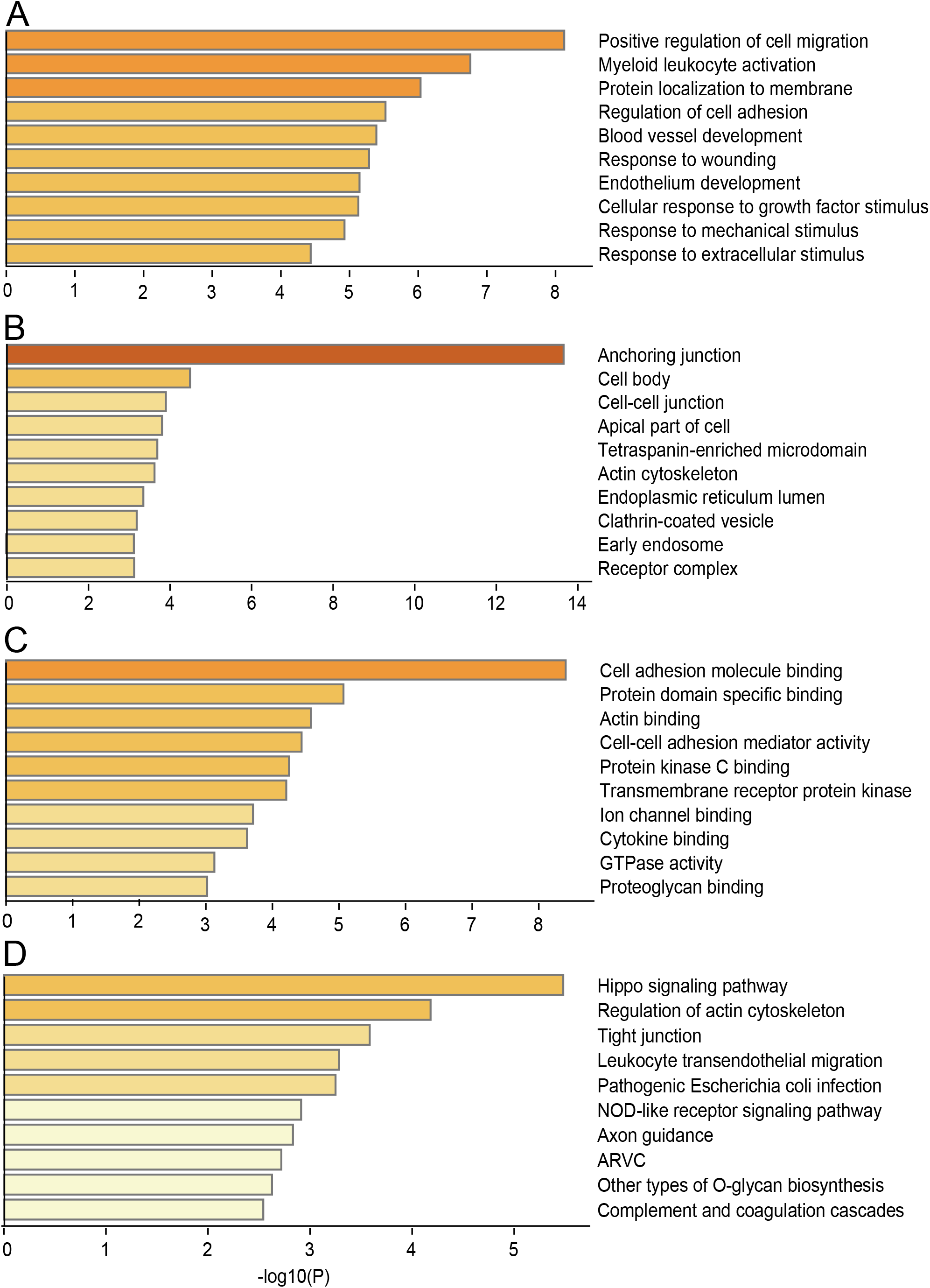
The top 10 significant GO terms of the enriched (A) biological process (B) Cell component (C) molecular function and (D) KEGG pathway.

**Figure 2.**
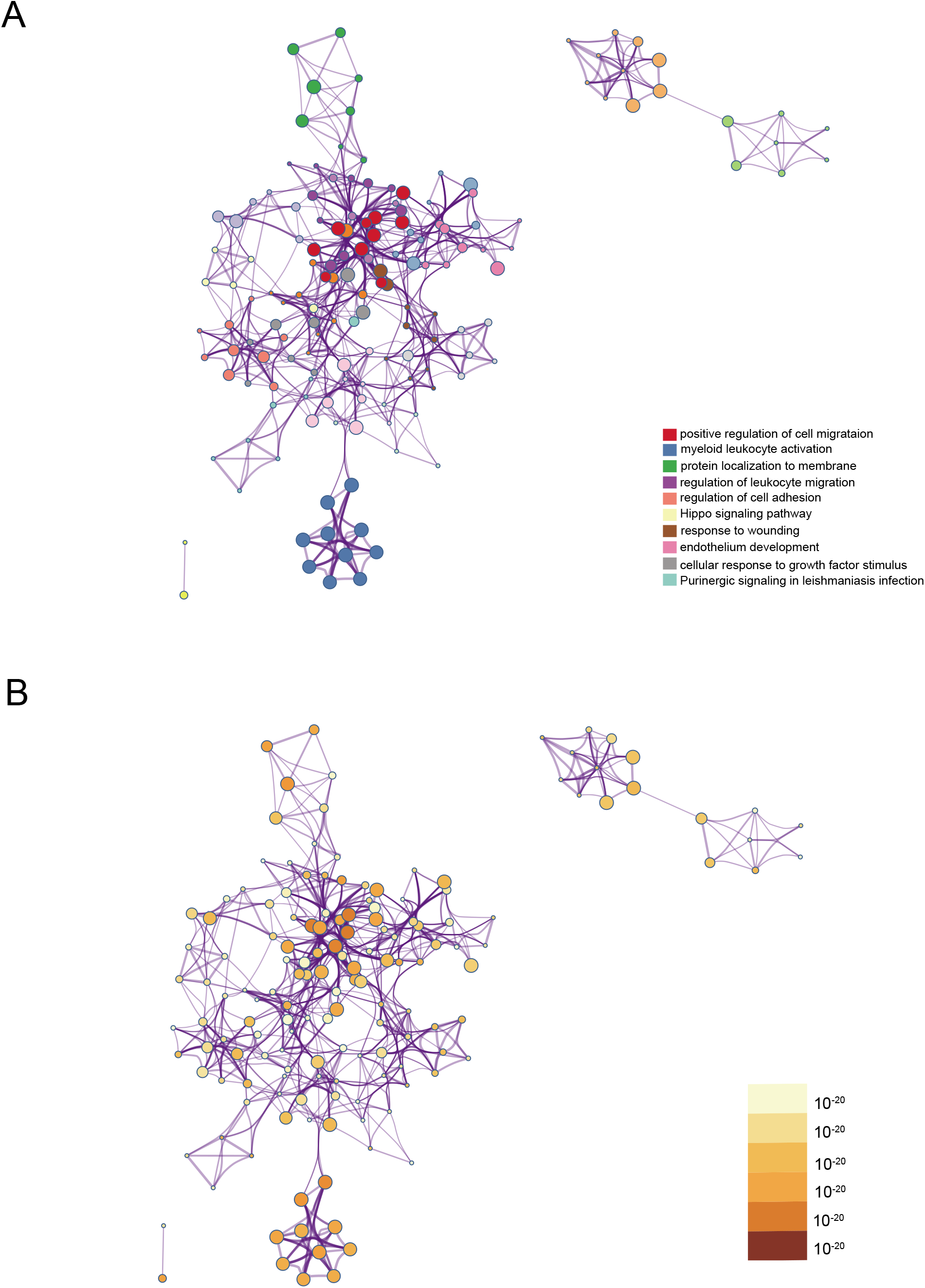
Network of enriched terms: (A) colored by cluster ID, where nodes that share the same cluster ID are typically close to each other; (B) colored by p-value, where terms containing more genes tend to have a more significant p-value.

### 3.3 The PPI of the DEGs

The PPI network is shown in Figure 3. There are 293 nodes and 514 edges included in this network, which accounted for 83.5% of the differentially expressed genes. Upregulated genes are represented in red color, while downregulated genes are represented in green color.

**Figure 3.**
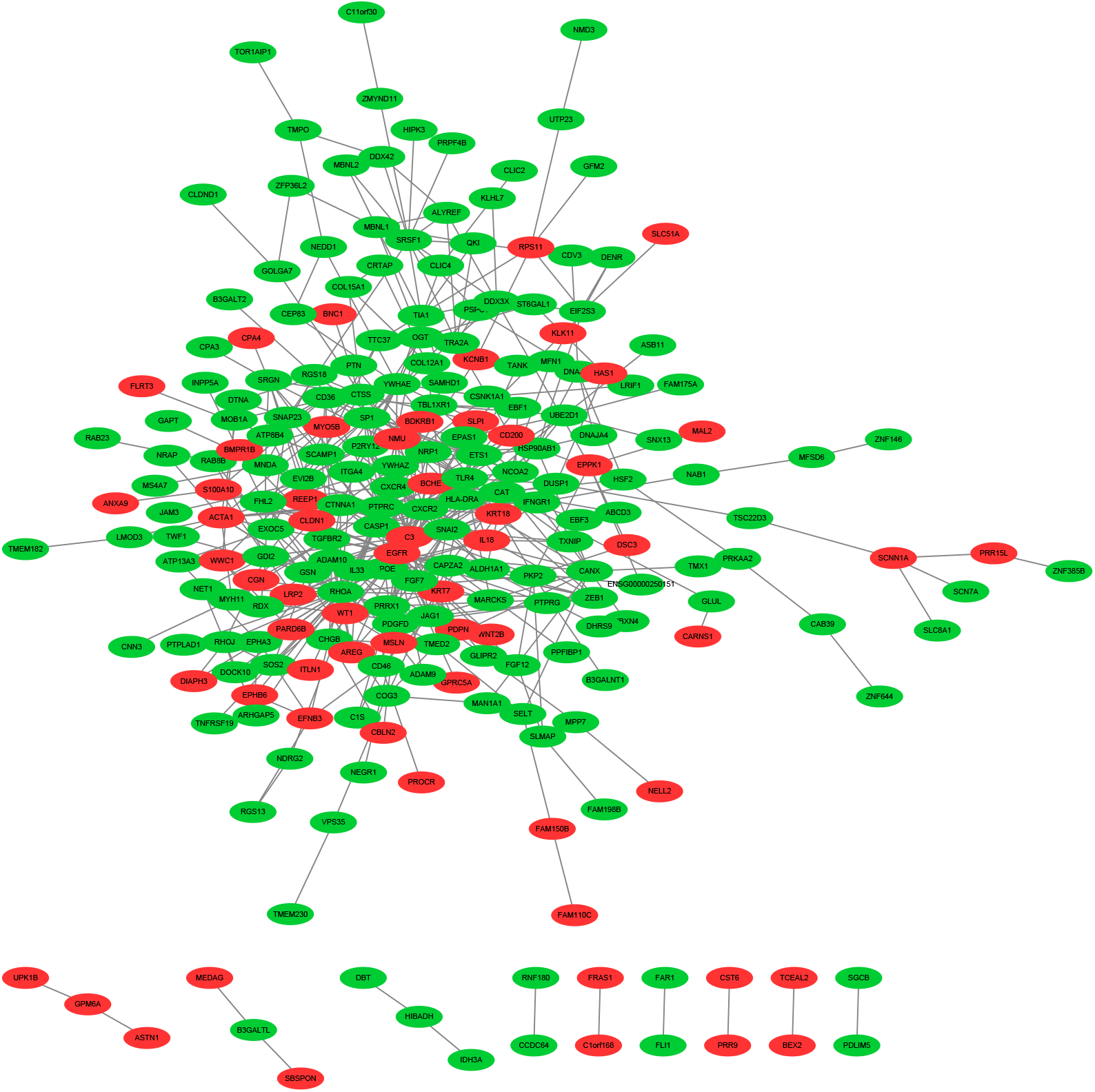
PPI network of DEGs. Upregulated genes were colored red, downregulated genes were colored green.

### 3.4 Module

To investigate densely connected network components, an MCODE network was constructed (Figure 4 and Table 1). The three best-scoring terms by p-value are shown in Figure and Table. A functional description of the corresponding components is also shown in the figure. The GO description of MCODE1 is Class A/1 (rhodopsin-like receptors), G alpha (i) signaling events, and peptide ligand-binding receptors. The GO description of MCODE2 is positive regulation of cellular protein localization, PI3K-Akt signaling pathway, and regulation of cellular protein localization. The GO description of MCODE3 is EPH-ephrin mediated repulsion of cells, ephrin receptor signaling pathway, and EPH-Ephrin signaling. GO description of MCODE4 is PID HIF2 PATHWAY, response to oxidative stress, Cellular responses to stress.

**Figure 4.**
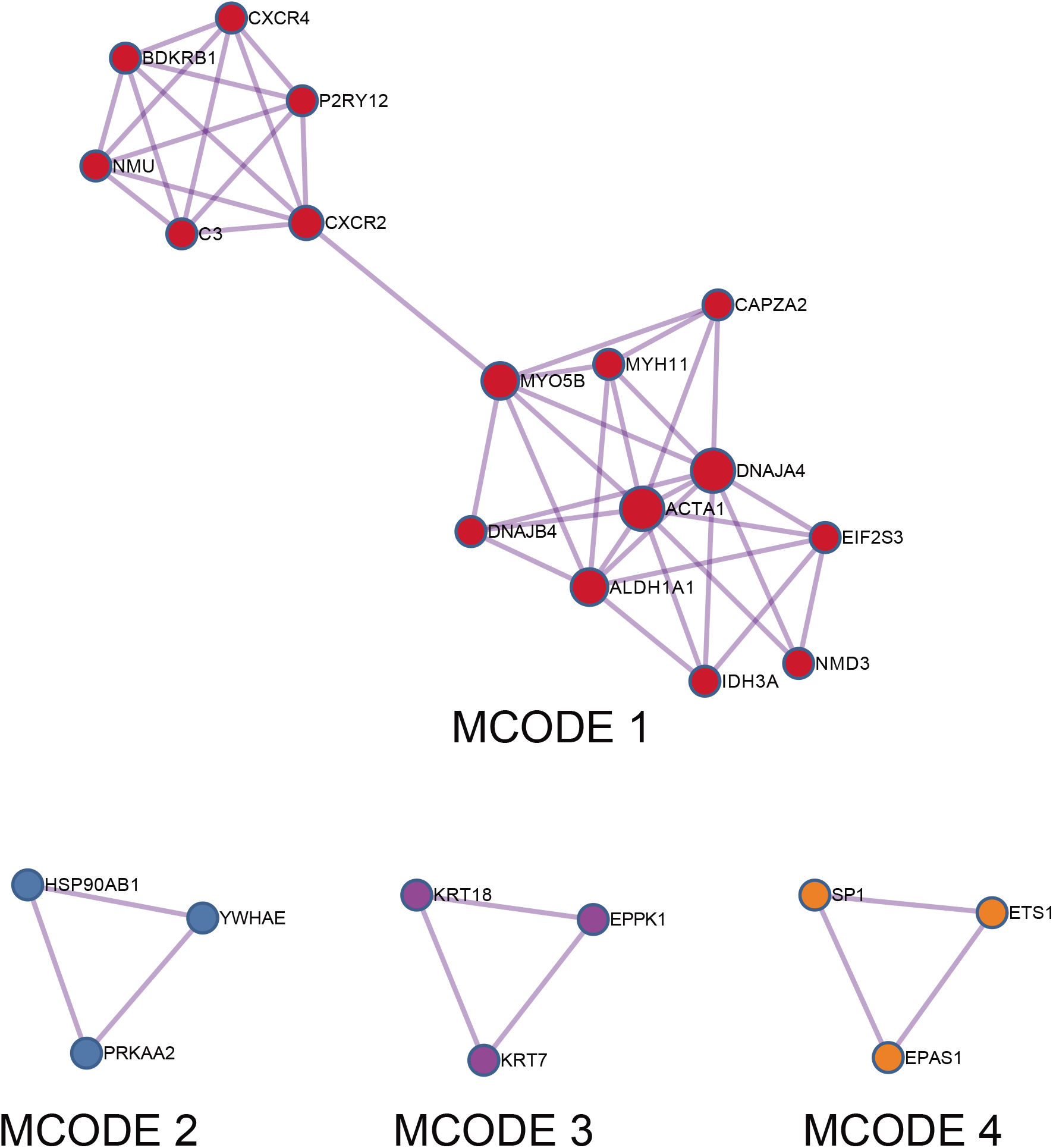
MCODE components identified in the gene lists.

### 3.5 Hub gene selection

The top 10 hub genes were determined by cytoHubba via the MCC method. The hub genes are shown in Figure 5. The upregulated genes were IL18 (degree=6), EGFR (degree=5), NMU (degree=4), and C3 (degree=4). The downregulated genes were CXCR2 (degree=9), TLR4 (degree=8), CXCR4 (degree=7), PTPRC (degree=7), CASP1 (degree=6), and IL33 (degree=6). Detailed information on these genes is shown in Table 2.

**Figure 5.**
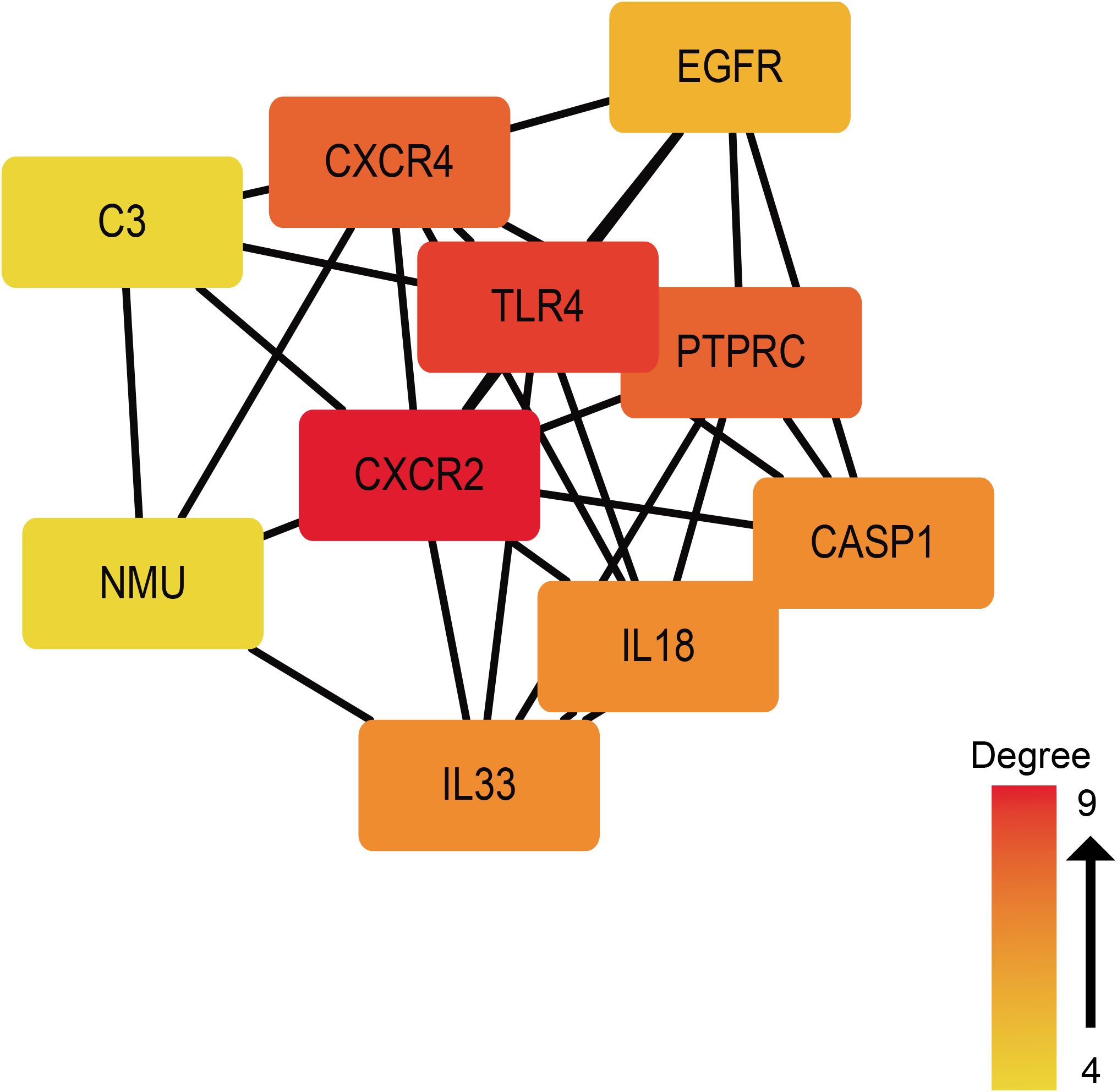
PPI network of 10 hub genes.

### 3.6 Analysis of TFs

We used PASTAA to predict transcription factors of hub genes. These transcription factors are shown in Table 3 and Table 4. The Irf-1 and Irf-10 families are the top transcription factors of the downregulated hub genes by p-value. For1 and For2 families are the top transcription factors of upregulated hub genes by p-value.

## Discussion

AF is the most common sustained arrhythmia in the world. Current therapies for AF are imperfect due to the incomplete understanding of the mechanism. Revealing the mechanism may help us treat AF better. Bioinformatic analysis of omics data is a powerful tool for mechanistic investigation. In our study, 93 upregulated and 268 downregulated genes were identified in patients with AF compared with patients with SR. The GO biological process enrichment analysis showed that positive cell migration was the most significantly enriched biological process. The regulation of cell migration has been proven to be related to atrial fibrillation occurrence and maintenance[16]. It has been reported that inhibition of fibroblast migration may attenuate atrial fibrosis and reduce AF vulnerability[17]. It has also been reported that inhibition of cell migration may prevent postinfarct atrial fibrillation in rats[18]. The anchoring junction was the most significantly enriched cellular component in the GO analysis. Growing evidence suggests that anchoring junctions play an important role in maintaining normal conduction. Disturbance of the anchoring junction has been proven to be related to arrhythmia, especially arrhythmogenic cardiomyopathy and Brugada syndrome[19, 20]. Abnormal propagation of action potential has been known to contribute to AF[21]. Anchoring junctions may be a potential target of AF treatment. In GO molecular function analysis, cell adhesion molecule binding was the most significantly enriched molecular function. Similar to cell migration, cell adhesion has also been reported to be related to atrial fibrillation[16].

In KEGG pathway analysis, the Hippo signaling pathway was the most enriched pathway. Studies have shown that Hippo signaling is related to arrhythmia. Xu’s research reported that Hippo signaling was likely to play a substantial role in the preventive effects of mechanical ventricular arrhythmia in response to left ventricle afterload increase[22]. Studies have also proven that Hippo signaling is implicated in arrhythmogenic cardiomyopathy pathogenesis and adipogenesis by regulating cell-cell contact[23]. Changes in cell-cell contact may affect the excitation and conduction of atrial cardiomyocytes, which may cause atrial arrhythmia, including atrial fibrillation. Therefore, we hypothesize that Hippo signaling may be involved in atrial fibrillation. Among the identified hub genes, CXCR2, TLR4 and CXCR4 were the top 3 genes. Recent studies have demonstrated that CXCR2 has been implicated in the pathogenesis of Ang-II-induced cardiac remodeling[24, 25]. Atrial remodeling, such as atrial fibrosis, contributes to the pathogenesis of atrial fibrillation. TLR4 has been reported to be related to new-onset atrial fibrillation in acute myocardial infarction[25]. Soppert’s study has shown that CXCR4 is involved in myofibroblast necroptosis, which may modulate cardiac remodeling in heart failure[26]. Based on these studies, we speculate that these 3 genes may be a novel target for atrial fibrillation treatment.

In modules extracted in the PPI network, Class A/1 (rhodopsin-like receptors) may be the critical module. It has high sequence similarity to the angiotensin receptor AT1 and plays an important role in the occurrence and development of cardiovascular and metabolic diseases, including atherosclerosis (AS), coronary heart disease (CAD), heart failure (HF), pulmonary arterial hypertension (PAH), myocardial hypertrophy and atrial fibrillation[27]. The hub genes CXCR2 and CXCR4 were included in this module, so the critical role of this module was further confirmed.

Transcription factors (TFs) may play important roles in gene expression. In the predicted TFs of downregulated genes, Irf-1 has been reported to be required in cardiac remodeling in response to pressure overload[28].

Our study may increase the understanding of atrial fibrillation mechanisms. Based on GO analysis, KEGG pathway analysis, the PPI network, hub gene prediction, key module prediction and transcription factor prediction, we found that the CXCR4, TLR4, and CXCR2 genes and the Class A/1 (rhodopsin-like receptors) module may play critical roles in atrial fibrosis. Hippo signaling may also contribute to atrial fibrillation occurrence.

Although our study may provide novel targets for atrial fibrillation treatment, there are still limitations in our study. The number of samples is relatively small, so there may be confounding factors. More samples are needed to confirm our findings. In addition to nuclear DNA, mitochondrial DNA (mtDNA) also carries genes. Mutation of mtDNA may also cause atrial fibrillation because elevated ROS levels have been proven to accelerate atrial fibrillation[29]. In our study, we did not analyze mtDNA data, and further studies on mtDNA are needed to determine the effect of mtDNA mutations in atrial fibrillation.

## Conclusion

Our study found that the CXCR4, TLR4, and CXCR2 genes and the class A/1 (rhodopsin-like receptors) module may play critical roles in atrial fibrosis. Hippo signaling may also contribute to atrial fibrillation occurrence. This may provide novel targets for AF treatment.

## Conflicts of interest

None.

## Notes

### Competing Interest Statement

The authors have declared no competing interest.

## Refenrence

1. Heijman, J., et al., Cellular and molecular electrophysiology of atrial fibrillation initiation, maintenance, and progression. Circ Res, 2014. 114 (9): p. 1483–99.

2. Zou, R., et al., Bioinformatic gene analysis for potential biomarkers and therapeutic targets of atrial fibrillation-related stroke. J Transl Med, 2019. 17(1): p. 45.

3. Morin, D.P., et al., The State of the Art: Atrial Fibrillation Epidemiology, Prevention, and Treatment. Mayo Clin Proc, 2016. 91(12): p. 1778–1810.

4. Dobrev, D. and S. Nattel, New antiarrhythmic drugs for treatment of atrial fibrillation. Lancet, 2010. 375(9721): p. 1212–23.

5. Dobrev, D., L. Carlsson, and S. Nattel, Novel molecular targets for atrial fibrillation therapy. Nat Rev Drug Discov, 2012. 11 (4): p. 275–91.

6. Heijman, J., N. Voigt, and D. Dobrev, New directions in antiarrhythmic drug therapy for atrial fibrillation. Future Cardiol, 2013. 9(1): p. 71–88.

7. Zhou, Y., et al., Metascape provides a biologist-oriented resource for the analysis of systems-level datasets. Nat Commun, 2019. 10(1): p. 1523.

8. Tsai, F.C., et al., Differential left-to-right atria gene expression ratio in human sinus rhythm and atrial fibrillation: Implications for arrhythmogenesis and thrombogenesis. Int J Cardiol, 2016. 222: p. 104–112.

9. Huang, H., et al., Identification of Potential Gene Interactions in Heart Failure Caused by Idiopathic Dilated Cardiomyopathy. Med Sci Monit, 2018. 24: p. 7697–7709.

10. Yu, G., et al., clusterProfiler: an R package for comparing biological themes among gene clusters. OMICS, 2012. 16(5): p. 284–7.

11. Reiner-Benaim, A., FDR control by the BH procedure for two-sided correlated tests with implications to gene expression data analysis. Biom J, 2007. 49(1): p. 107–26.

12. Huang da, W., B.T. Sherman, and R.A. Lempicki, Systematic and integrative analysis of large gene lists using DAVID bioinformatics resources. Nat Protoc, 2009. 4(1): p. 44–57.

13. Szklarczyk, D., et al., The STRING database in 2017: quality-controlled protein-protein association networks, made broadly accessible. Nucleic Acids Res, 2017. 45(D1): p. D362–D368.

14. Altaf-Ul-Amin, M., et al., Development and implementation of an algorithm for detection of protein complexes in large interaction networks. BMC Bioinformatics, 2006. 7: p. 207.

15. Min, F., F. Gao, and Z. Liu, Screening and further analyzing differentially expressed genes in acute idiopathic pulmonary fibrosis with DNA microarray. Eur Rev Med Pharmacol Sci, 2013. 17(20): p. 2784–90.

16. Wu, Y.X., et al., Time Series Gene Expression Profiling and Temporal Regulatory Pathway Analysis of Angiotensin II Induced Atrial Fibrillation in Mice. Front Physiol, 2019. 10: p. 597.

17. Song, S., et al., EZH2 as a novel therapeutic target for atrial fibrosis and atrial fibrillation. J Mol Cell Cardiol, 2019. 135: p. 119–133.

18. Ma, J., et al., Matrine reduces susceptibility to postinfarct atrial fibrillation in rats due to antifibrotic properties. J Cardiovasc Electrophysiol, 2018. 29(4): p. 616–627.

19. Moncayo-Arlandi, J. and R. Brugada, Unmasking the molecular link between arrhythmogenic cardiomyopathy and Brugada syndrome. Nat Rev Cardiol, 2017. 14(12): p. 744–756.

20. Vermij, S.H., H. Abriel, and T.A. van Veen, Refining the molecular organization of the cardiac intercalated disc. Cardiovasc Res, 2017. 113 (3): p. 259–275.

21. Staerk, L., et al., Atrial Fibrillation: Epidemiology, Pathophysiology, and Clinical Outcomes. Circ Res, 2017. 120(9): p. 1501–1517.

22. Xu, X., et al., Effects of artemisinin on ventricular arrhythmias in response to left ventricular afterload increase and microRNA expression profiles in Wistar rats. PeerJ, 2018. 6: p. e6110.

23. Austin, K.M., et al., Molecular mechanisms of arrhythmogenic cardiomyopathy. Nat Rev Cardiol, 2019. 16(9): p. 519–537.

24. Wang, L., et al., CXCL1-CXCR2 axis mediates angiotensin II-induced cardiac hypertrophy and remodelling through regulation of monocyte infiltration. Eur Heart J, 2018. 39(20): p. 1818–1831.

25. Zhang, P., L. Shao, and J. Ma, Toll-Like Receptors 2 and 4 Predict New-Onset Atrial Fibrillation in Acute Myocardial Infarction Patients. Int Heart J, 2018. 59(1): p. 64–70.

26. Soppert, J., et al., Soluble CD74 Reroutes MIF/CXCR4/AKT-Mediated Survival of Cardiac Myofibroblasts to Necroptosis. J Am Heart Assoc, 2018. 7(17): p. e009384.

27. Cao, J., H. Li, and L. Chen, Targeting drugs to APJ receptor: the prospect of treatment of hypertension and other cardiovascular diseases. Curr Drug Targets, 2015. 16(2): p. 148–55.

28. Jiang, D.S., et al., Interferon regulatory factor 1 is required for cardiac remodeling in response to pressure overload. Hypertension, 2014. 64(1): p. 77–86.

29. Li, X., et al., Mitochondria and the Pathophysiological Mechanism of Atrial Fibrillation. Curr Pharm Des, 2018. 24(26): p. 3055–3061.

