## Supplementary material for "Bioinformatic analysis for the identification of potential gene interactions and therapeutic targets in atrial fibrillation": Table 1

| **MCODE** | **GO** | **Description** | **Log10(P)** |
| --- | --- | --- | --- |
| MCODE_1 | R-HSA-373076 | Class A/1 (Rhodopsin-like receptors) | -7.3 |
| MCODE_1 | R-HSA-418594 | G alpha (i) signalling events | -6.8 |
| MCODE_1 | R-HSA-375276 | Peptide ligand-binding receptors | -6.8 |
| MCODE_2 | GO:1903829 | positive regulation of cellular protein localization | -5.6 |
| MCODE_2 | hsa04151 | PI3K-Akt signaling pathway | -5.6 |
| MCODE_2 | GO:1903827 | regulation of cellular protein localization | -5 |
| MCODE_3 | R-HSA-3928665 | EPH-ephrin mediated repulsion of cells | -8.1 |
| MCODE_3 | GO:0048013 | ephrin receptor signaling pathway | -7.4 |
| MCODE_3 | R-HSA-2682334 | EPH-Ephrin signaling | -7.3 |
| MCODE_4 | M44 | PID HIF2PATHWAY | -8.6 |
| MCODE_4 | GO:0006979 | response to oxidative stress | -5.2 |
| MCODE_4 | R-HSA-2262752 | Cellular responses to stress | -4.8 |

**Table 1. Pathway and process enrichment analysis**

GO: Gene ontology
