## Supplementary material for "Bioinformatic analysis for the identification of potential gene interactions and therapeutic targets in atrial fibrillation": Table 2

| **Gene** | **LogFC** | **P** | **Degree** |
| --- | --- | --- | --- |
| CXCR2 | -1.50 | 0.022 | 9 |
| TLR4 | -1.36 | 0.027 | 8 |
| CXCR4 | -1.30 | 0.030 | 7 |
| PTPRC | -1.26 | 0.017 | 7 |
| CASP1 | -1.10 | 0.015 | 6 |
| IL33 | -1.07 | 0.020 | 6 |
| IL18 | 1.39 | 0.014 | 6 |
| EGFR | 1.15 | 0.001 | 5 |
| NMU | 1.79 | 0.048 | 4 |
| C3 | 1.20 | 0.026 | 4 |

**Table 2 Description of 10 hub genes**
