## Supplementary material for "Bioinformatic analysis for the identification of potential gene interactions and therapeutic targets in atrial fibrillation": Table 3

| **Rank** | **Matrix** | **Transcription Factor** | **Association Score** | **P-Value** |
| --- | --- | --- | --- | --- |
| 1 | IRF_Q6 | Irf-1, Irf-10 | 4.212 | 2.04E-04 |
| 2 | IRF_Q6_01 | Irf-1, Irf-10 | 3.366 | 9.41E-04 |
| 3 | PU1_Q6 | Pu.1 | 3.123 | 1.68E-03 |
| 4 | ETS_Q6 | Elf-1, Elfr | 3.106 | 1.68E-03 |
| 5 | ICSBP_Q6 | [Irf-8](http://www.genecards.org/cgi-bin/carddisp.pl?gene=Irf-8) | 3.015 | 2.88E-03 |
| 6 | CEBP_Q2_01 | C/ebpalpha , C/ebpalpha(p30) | 2.72 | 5.95E-03 |
| 7 | PEA3_Q6 | [Pea3](http://www.genecards.org/cgi-bin/carddisp.pl?gene=Pea3) | 2.646 | 6.45E-03 |
| 8 | TATA_C | [Tbp](http://www.genecards.org/cgi-bin/carddisp.pl?gene=Tbp) | 2.574 | 6.83E-03 |
| 9 | SEF1_C | N/A | 2.522 | 9.05E-03 |
| 10 | INR_HAND100 | N/A | 2.473 | 9.47E-03 |
| 11 | SRF_Q5_01 | [Srf](http://www.genecards.org/cgi-bin/carddisp.pl?gene=Srf) | 2.346 | 1.31E-02 |
| 12 | CEBP_Q3 | C/ebp, C/ebpalpha | 2.282 | 1.44E-02 |
| 13 | BLIMP1_Q6 | [Blimp-1](http://www.genecards.org/cgi-bin/carddisp.pl?gene=Blimp-1) | 2.278 | 1.44E-02 |
| 14 | OCT4_01 | N/A | 2.221 | 1.70E-02 |
| 15 | POLY_C | N/A | 2.221 | 1.70E-02 |
| 16 | COUPTF_Q6 | Coup, Coup-tf1 | 2.221 | 1.70E-02 |
| 17 | HNF4_Q6 | Hnf-4, Hnf-4alpha | 2.123 | 2.10E-02 |
| 18 | MEIS1AHOXA9_01 | Hoxa9, Hoxa9b | 2.046 | 2.25E-02 |
| 19 | MEIS1BHOXA9_02 | Hoxa9, Hoxa9b | 2.046 | 2.25E-02 |
| 20 | OCT4_02 | N/A | 2.046 | 2.25E-02 |

**Table 3. Top 20 transcription factors of down-regulated genes**
