## Supplementary material for "Bioinformatic analysis for the identification of potential gene interactions and therapeutic targets in atrial fibrillation": Table 4

| **Rank** | **Matrix** | **Transcription Factor** | **Association Score** | **P-Value** |
| --- | --- | --- | --- | --- |
| 1 | FXR_IR1_Q6 | For1, For2 | 2.965 | 3.15E-03 |
| 2 | AP4_01 | [Ap-4](http://www.genecards.org/cgi-bin/carddisp.pl?gene=Ap-4) | 2.876 | 3.52E-03 |
| 3 | AP2ALPHA_03 | N/A | 2.346 | 1.31E-02 |
| 4 | R_01 | N/A | 2.346 | 1.31E-02 |
| 5 | CEBPB_01 | C/ebpbeta(lap), C/ebpbeta(p35) | 2.221 | 1.70E-02 |
| 6 | DPE_HAND | N/A | 2.221 | 1.70E-02 |
| 7 | PAX4_01 | [Pax-4a](http://www.genecards.org/cgi-bin/carddisp.pl?gene=Pax-4a) | 1.976 | 2.50E-02 |
| 8 | BARBIE_01 | N/A | 1.919 | 2.91E-02 |
| 9 | DR3_Q4 | Car, Pxr-1 | 1.87 | 3.16E-02 |
| 10 | E2F1_Q4 | [E2f-1](http://www.genecards.org/cgi-bin/carddisp.pl?gene=E2f-1) | 1.868 | 3.16E-02 |
| 11 | AP2ALPHA_02 | [Ap-2alphaa](http://www.genecards.org/cgi-bin/carddisp.pl?gene=Ap-2alphaa) | 1.824 | 3.41E-02 |
| 12 | MTATA_B | N/A | 1.743 | 3.84E-02 |
| 13 | AP2_Q3 | Ap-2alpha, Ap-2alphaa | 1.618 | 5.01E-02 |
| 14 | SP1_01 | [Sp1](http://www.genecards.org/cgi-bin/carddisp.pl?gene=Sp1) | 1.617 | 5.01E-02 |
| 15 | SF1_Q6_01 | N/A | 1.606 | 5.01E-02 |
| 16 | E2_Q6_01 | N/A | 1.57 | 5.58E-02 |
| 17 | TFIII_Q6 | [Tfii-i](http://www.genecards.org/cgi-bin/carddisp.pl?gene=Tfii-i) | 1.57 | 5.58E-02 |
| 18 | TTF1_Q6 | [Nkx2-1](http://www.genecards.org/cgi-bin/carddisp.pl?gene=Nkx2-1) | 1.567 | 5.58E-02 |
| 19 | ETF_Q6 | N/A | 1.446 | 6.98E-02 |
| 20 | SPZ1_01 | [Spz1](http://www.genecards.org/cgi-bin/carddisp.pl?gene=Spz1) | 1.446 | 6.98E-02 |

**Table 4. Top 20 transcription of up-regulated genes**
